# Why is the perceptual octave stretched? An account based on mismatched time constants within the auditory brainstem

**DOI:** 10.1101/2022.12.05.519114

**Authors:** Alain de Cheveigné

## Abstract

This paper suggests an explanation for listeners’ greater tolerance to positive than negative mistuning of the higher tone within an octave pair. It hypothesizes a neural circuit tuned to cancel the lower tone, that also cancels the higher tone if that tone is in tune. Imperfect cancellation is the cue to mistuning of the octave. The circuit involves two neural pathways, one delayed with respect to the other, that feed a coincidence-sensitive neuron via excitatory and inhibitory synapses. A mismatch between the time constants of these two synapses results in an asymmetry in sensitivity to mismatch. Specifically, if the time constant of the *delayed* pathway is greater than that of the direct pathway, there is a greater tolerance to *positive* mistuning than to negative mistuning. The model is directly applicable to the harmonic octave (concurrent tones), but extending it to the melodic octave (successive tones) requires additional assumptions that are discussed. The paper reviews evidence from auditory psychophysics and physiology in favor – or against – this explanation.

## I. INTRODUCTION

It sometimes happens that competing theories differ on subtle effects that allow us to decide between them. A famous example is the precession of the perihelion of Mercury, a tiny effect (43 arc-seconds per century) that provided support for General Relativity over Newtonian physics. The asymmetry of mistuning for the octave interval (Demany et al., 2021, 1991) might play a similar role in hearing.

The octave (frequency ratio 2:1) is one of a series of small integer ratios that appear naturally between partials of a harmonic complex sound, or between the subharmonics of any periodic tone, complex or pure. Tones with frequencies in this ratio display a perceptual affinity when they are played together (harmonic octave) or in succession (melodic octave). The octave plays a special role in music of many cultures and genres, and there is some evidence that melodic octave similarity (if not “equivalence”) is innate (Demany and Armand, 1984). Listeners, particularly musicians, are sensitive to small mistunings from the exact octave. “Asymmetry” refers to the fact that a compressed octave (ratio smaller than 2) is better detected than a stretched octave (ratio larger than 2).

Explanations of octave affinity have been imagined for both spectral and temporal theories of auditory processing. For example Terhardt (1974) proposed that listeners acquire harmonic templates from exposure to harmonic sounds, whereas Ohgushi (1983) remarked that intervals between auditory nerve spikes (action potentials) evoked by the lower tone are also found between spikes evoked by the octave. Both ideas can be developed into models to account for harmonic and melodic octave affinity (Demany et al., 2021).

To the extent that those explanations involve integer ratios, it seems odd that the octave might be perceptually stretched. Deciding if the pitches of two *consecutive* pure tones form an octave requires enough musical skill to at least understand instructions, and a common outcome is that the frequency of the upper tone (*f_H_*) is matched higher than twice the lower (2*f_L_*) by a few percent (Ward, 1954). This is known as melodic “octave stretching” or “enlargement”. Mistuning from an octave relationship between two *concurrent* pure tones can likewise be detected. Here, it is observed that tolerance is greater for positive than negative mistuning (Demany et al., 1991). It is natural to search for a common explanation for melodic stretch and harmonic mistuning asymmetry, but experimental support has been elusive, some studies favoring distinct mechanisms (Bonnard et al., 2016, 2013), others a common mechanism (Demany et al., 2021).

The aim of this paper is to explore an explanation of mistuning asymmetry according to which the asymmetry arises from a mismatch in time constants of neural signal processing networks within the auditory brainstem. This explanation applies to the harmonic octave, but extending it to the melodic octave requires additional assumptions that are considered. The psychophysics of octave tuning are further reviewed in the Discussion, together with relevant evidence from auditory physiology.

## II. METHODS AND RESULTS

This section introduces the required concepts and tools, and describes the main result.

### a. Stimuli

In contrast to the harmonics-rich sounds of many musical instruments, most studies employ pure tones, so as to avoid the confound of beats between partials. A pure tone of frequency *f* can, however, be treated as a harmonic complex with only one harmonic, with “fundamental frequency” *f* and period *T* = 1/*f*.

### b. Cochlear frequency analysis

As a first step of auditory processing, stimuli undergo filtering in the cochlea, modeled here as a gammatone filter bank with bandwidths from Moore and Glasberg (1983). Those bandwiths are roughly 1/3 octave (except in the low frequency region where filters are wider), so pure tones spaced one octave apart are well resolved, as illustrated in Fig. 1. Channels with characteristic frequencies (CFs) close to a tone’s frequency are dominated by that tone by up to 50 dB or more, whereas some intermediate channels respond to both.

**FIG. 1.**
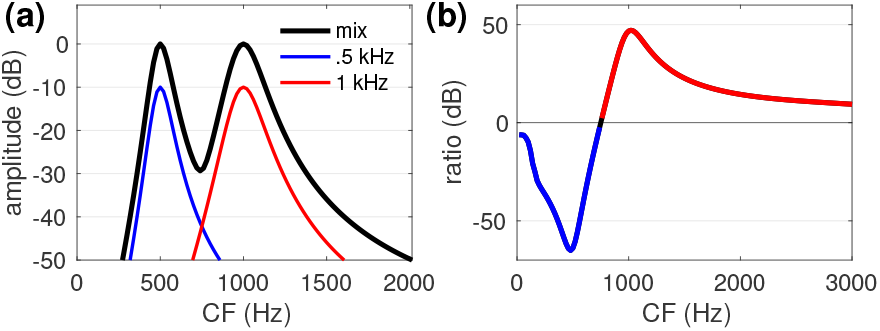
(a) Power as a function of filter center frequency (CF) for a pair of pure tones separated by an octave (black) and for each tone alone (blue and red, offset vertically for clarity). (b) Power of higher tone divided by power of lower tone (blue: lower dominates, red: upper dominates). These patterns were derived from a gammatone filterbank implementation (Slaney, 1993) with bandwidth parameters from Moore and Glasberg (1983).

Figure 1 gives us an idea of the information available for the auditory brain to detect the octave relationship between two pure tones. The *profile of excitation across filter CF* (panel a) shows a clear peak at the frequencies of both tones, which could thus be extracted from a tonotopic pattern. Multiple channels are dominated by one tone or the other (panel b), so information about those frequencies is also available from the *temporal fine structure* of auditory nerve discharges. Finally, a smaller set of intermediate channels responds to both tones (circa 700 Hz) possibly offering a third temporal cue in the form of their interaction. It is worth noting that these patterns depend largely on the shape of the *skirts* of cochlear filter transfer functions, whereas most filter models are optimized for their shape near the peak (Patterson, 1976).

### c. Period estimation and pitch

A popular model of pitch perception suggests that pitch is derived from the temporal fine structure of auditory nerve discharges, for example according to an autocorrelation-like mechanism (Cariani and Delgutte, 1996; Licklider, 1951; Meddis and Hewitt, 1991). Using a sampled-signal notation, the autocorrelation function (ACF) can be defined as:

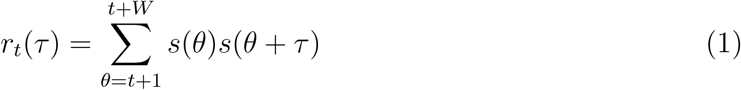

where *s*(*t*) is the acoustic signal (or some transform of it), *W* is an integration window, *τ* is the time at which the function is calculated, and *τ* is the lag parameter (Fig. 2, left). Applied to a periodic stimulus, such as a pure tone, the function shows a peak at all multiples of *T*, where *T* = 1/*f*_0_ is the period. An alternative to the ACF is the the “Squared Difference Function” (SDF):

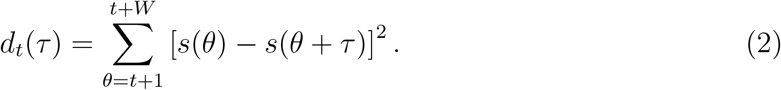

calculated as the Euclidean distance between two windows, one fixed the other shifted by *τ*. These functions are closely related: *d_t_*(*τ*) = *p_t_*(*τ*) – 2*r_t_*(*τ*), where *p_t_*(*τ*) = *r_t_*(0) + *r_t+τ_*(0) reflects signal power circa time *t*. Applied to a periodic stimulus, such as pure tone, the SDF takes its smallest possible value (zero) at *T* and all its multiples (Fig. 2, right).

**FIG. 2.**
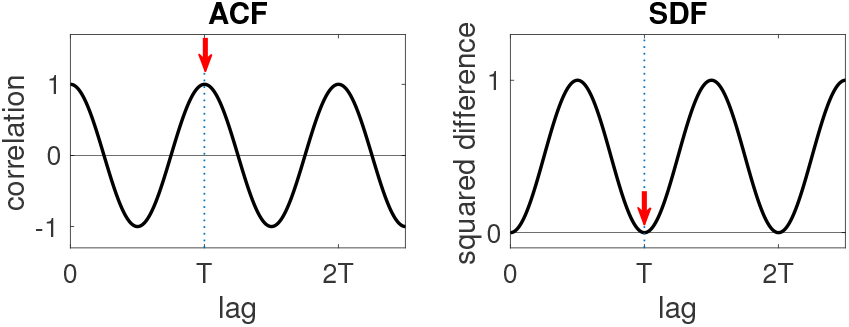
Period estimation based on autocorrelation (left) or cancellation (right). For a periodic stimulus of period *T*, the autocorrelation function (ACF) shows a peak for a lag equal to *T* (left), while the squared difference function (SDF) shows a zero at that same lag (right). Both can be used as a cue to the period (and thus the pitch) of the stimulus. The functions here are normalized to take values between −1 and 1 (ACF) or 0 and 1 (SDF).

In the *autocorrelation model of pitch*, a neural approximation of the ACF is calculated within every channel of a peripheral filter bank (Licklider, 1951). In response to a stimulus of period *T*, all channels show a peak at *T*, and the position of this common peak is the cue to the period of a sound, and thus its pitch. The CF × lag pattern can also be summarized by averaging it over all channels to form a *summary autocorrelation function* (SACF), that typically also shows a peak at *T* (Meddis and Hewitt, 1991). Alternatively, in some tasks it might be beneficial to restrict the pattern to a subset of channels. For example, looking at Fig. 1, it would seem best to estimate the period of the lower tone from channels with CF below ~700 Hz, and the period of the higher tone from channels with CF above that limit.

Replacing the ACF by the SDF yields a pitch model with properties very similar to the autocorrelation model, the cue to the period being the position of a dip (rather than a peak) in the CF × lag pattern (Fig. 2) (de Cheveigné, 1998). A reason for considering this variant is that it can easily be extended to detect mistuning.

### d. Cancellation and mistuning

The SDF can be understood as quantifying variance of the output of a time-domain “harmonic cancellation” filter

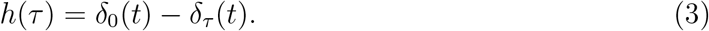

calculated over a window *W*, as a function of the delay parameter *τ*, where *δ_τ_* is the Kronecker delta function translated to *τ*. Figure 3 (left) shows such a filter with lag parameter *T*, together with its transfer function (Fig. 3 right). The transfer function of this filter shows deep dips at 1/*T* and its multiples, the output being zero for any periodic input signal with period *T/k*. Mistuning of a signal of frequency *f* away from *k/T* can be detected as an increase in power at the output of the filter.

**FIG. 3.**
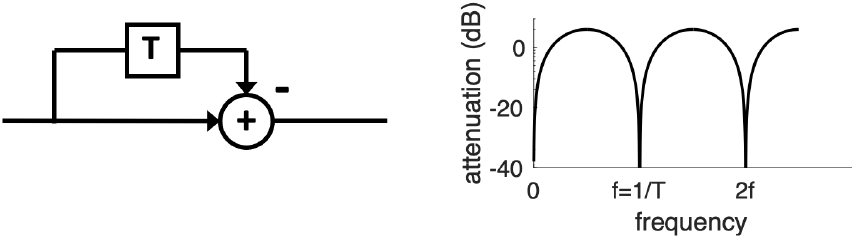
Harmonic cancellation filter. Left: comb filter with impulse response *δ*_0_(*t*) – *δ_T_*(*t*). Right: transfer function. This filter suppresses the fundamental and all harmonics of a stimulus with period *T*.

More generally, *harmonic cancellation* is a hypothetical process according to which the auditory system exploits harmonicity of a background sound to suppress it (de Cheveigné, 2021), for example using a harmonic cancellation filter (Eqn. 3). This process might explain why mistuning a component within a harmonic complex makes it more salient (Hartmann et al., 1990; Moore et al., 1985) and its pitch easier to match (Hartmann and Doty, 1996). Octave-spaced tones are a special case of this situation.

### e. Octave mistuning

Harmonic octave mistuning can be detected with the following steps: (1) estimate the period of the lower tone, (2) use it as the parameter of a cancellation filter applied to the higher tone, (3) decide that the octave is mistuned if the cancellation residual exceeds a threshold, and in-tune otherwise.

Referring to Fig. 2 (right), it appears that low CF channels represent well the lower tone, high CF channels the higher tone, and a smaller set of intermediate channels represent both. There are at least three plausible scenarios. A first is that the period is estimated in low-CF channels and used to parametrize a harmonic cancellation filter applied to high-CF channels. A second is that both operations are applied to the set of intermediate channels (avoiding the need to carry the estimate across channels). A third is that both are applied after cross-channel convergence (see section on physiology in the Discussion).

### f. Neural implementation

Processing within the auditory brain must rely on the spike-based neural representation created by transduction in the cochlea. Licklider (1951) imagined that the ACF might be approximated by an array of elementary neural circuits, each comprising a neuron fed from direct and delayed pathways via two excitatory synapses (Fig. 4, top left). The neuron fires only if spikes arrive together (within some short window) on both pathways. For example, short excitatory post synaptic potentials (EPSPs) evoked by spikes might interact additively (van der Heijden et al., 2013), reaching threshold only if they overlap in time (Fig. 4, top right). Instantaneous output spike probability then approximates the *product* of input spike probabilities, as required to approximate Eq. 1. Such a mechanism has been hypothesized within the medial superior olive (MSO) in a circuit that estimates interaural time differences (Lu et al., 2018; van der Heijden et al., 2013).

**FIG. 4.**
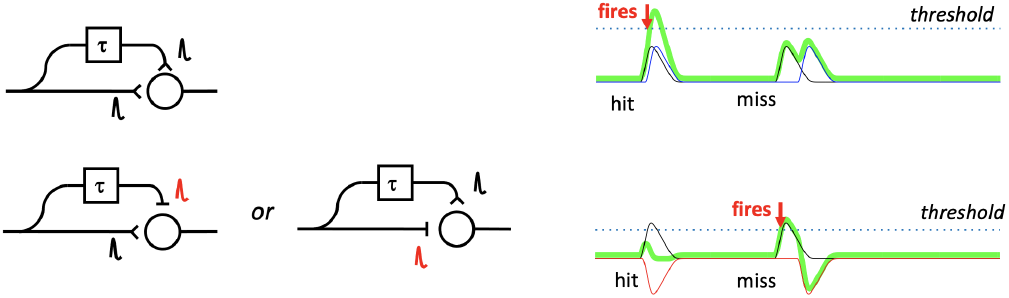
Neural approximations of autocorrelation (top) or cancellation (bottom). Top: a neuron receives direct and delayed input via two excitatory synapses and fires when spikes arrive simultaneously on both. This can be used to approximate the delay-and-multiply operation of the ACF. Bottom: a neuron receives direct and delayed input via excitatory and inhibitory synapses and fires whenever a spike arrives on the excitatory input, unless a spike arrives simultaneously on the inhibitory input. This can be used to approximate the delay-and-subtract operation of the harmonic cancellation filter, or the SDF. The delay can be applied indifferently to the inhibitory input (left) or to the excitatory input (center).

An analogous circuit can approximate the harmonic cancellation filter (de Cheveigné, 2021). A neuron is contacted by an excitatory and an inhibitory synapse, one fed from a direct pathway, the other via a delay (Fig. 4, bottom left). The EPSP and IPSP (inhibitory post synaptic potential) summate as in Fig. 4 (bottom right). The neuron’s threshold is such that it fires at each excitatory spike, *unless* a spike arrives together at the inhibitory synapse within some short window. Instantaneous output spike probability then reflects the difference of input spike probabilities, as required to approximate Eq. 2. Such a gating mechanism has been found within the lateral superior olive (LSO) in a circuit that estimates interaural differences in level or onset time (Franken et al., 2018).

### g. A neural octave mistuning detector

The neural cancellation mechanism can be used to detect octave mistuning. To be concrete, let us suppose that the lower tone has a frequency of 500 Hz. A harmonic cancellation filter is implemented as in Fig. 4 (left or center) with EPSP and IPSP shaped as in Fig. 5 (top left), with the inhibitory pathway delayed by *T* = 2 ms (inverse of 500 Hz). The spike train is split in two pathways, one convolved with the EPSP, and the other delayed and convolved with the IPSP (Fig. 4 bottom left). The firing probability, modelled as one minus the normalized cross-product between direct and delayed PSPs, is plotted in Fig. 5 (top right) as a function of frequency circa 1 kHz. It is minimal at 1 kHz (in tune) and increases on either side (mistuned). The increase in firing probability with mistuning would offer a cue to mistuning from the octave relationship.

**FIG. 5.**
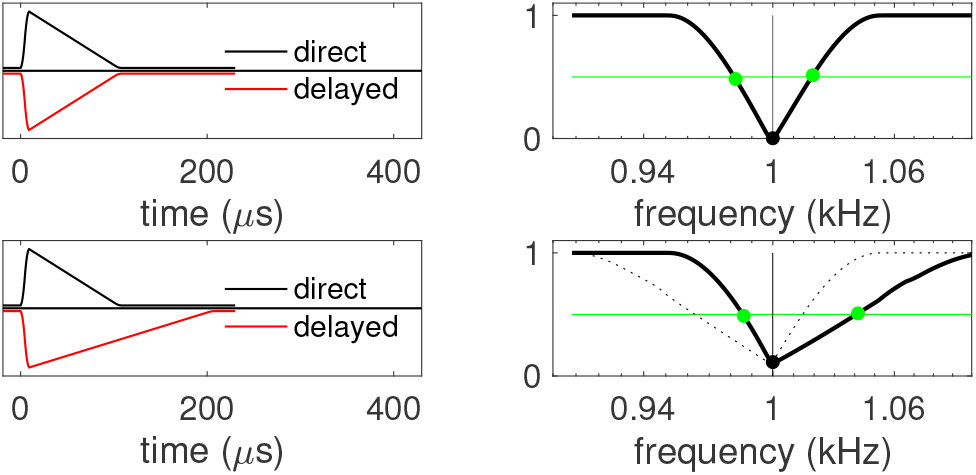
Detection of mistuning with a neural cancellation filter. Left: schematic PSPs on direct and delayed pathways (direct: EPSP, black, delayed: IPSP, red). Right: output of filter as a function of frequency of the higher tone in a mistuned-harmonic pair. Top: EPSP and IPSP follow similar dynamics, the mistuning curve is roughly symmetrical around 1 kHz. Bottom: the PSP on the delayed pathway is longer than that on the direct pathway. The mistuning curve is shallower for positive than for negative mistunings (green dots indicate the mistuning for which the filter output exceeds the threshold indicated by the green line). If the delayed PSP were the shorter one, the asymmetry would be reversed (dotted line). In all cases, the dip is centered precisely at the in-tune octave (1 kHz).

### h. Asymmetry

We are now in position to suggest an explanation for the mistuning asymmetry. First, let us suppose that PSPs on direct and delayed pathways have a similar duration: the mistuning curve is roughly symmetrical around 1 kHz (Fig. 5 top right), with slopes that depend on PSP duration, being shallower for longer PSPs. If instead the PSPs differ in duration, the mistuning curve is *asymmetrical* relative to 1 kHz. In particular, if the PSP duration of the delayed pathway is longer than that of the direct pathway, there is a greater tolerance for a positive mistuning (stretched octave) than negative. A threshold value of filter output (green dots) is reached for a smaller value of negative than positive mistuning, qualitatively consistent with most behavioral results. Interestingly, the position of the minimum does not shift, just the slopes.

The origin of the asymmetry can be understood intuitively by looking at Fig. 5 (left, bottom). Increasing the higher-tone period by 100 *μs* (compressed octave) would shift the onset of the delayed IPSP beyond the offset of the direct EPSP, abolishing the inhibitory effect. In constrast, decreasing it by the same amount (stretched octave) would only partially reduce the overlap. If, instead, the delayed path had the shorter PSP, the asymmetry would be reversed (dotted line in Fig. 5 bottom right). The asymmetry is also contingent on the asymmetric shape of the PSPs (steeper onset than offset).

The demonstration that asymmetry may arise within a plausible mechanism of octave mistuning detection is the main result of this paper. However, the predicted *direction* depends on time constants for delayed and undelayed pathways. To match experimental results, the delayed pathway must have a longer time constant than the direct pathway, if not, the model fails. The model is applicable directly to the harmonic octave (concurrent sounds); the degree to which it accounts also for melodic octave affinity is examined in the Discussion.

### i. Simulation

The model was simulated and its predictions compared with behavioral data from Demany et al. (1991). Stimuli in that study consisted of the sum of two octave-spaced pure tones of duration 0.5 s with equal amplitudes and lower tone frequency 300 Hz, 500 Hz, 1000 Hz or 2000 Hz. Here, the stimulus is fed to a gammatone filter bank (Slaney, 1993) with bandwidth parameters from Moore and Glasberg (1983). A single channel was selected, with CF equal to the frequency of the higher tone, and the half-wave rectified channel output scaled to drive an inhomogenous Poisson process with a 1 ms dead time (refractory effect). That was implemented as a homogenous Poisson process followed by a thinning process to enforce a time-dependent instantaneous rate and an interval-dependent refractory function (de Cheveigné, 1985; Lewis, 1979). The scaling factor was adjusted to ensure a physiologically-realistic average discharge rate *R*. To simulate the group of *N* auditory fibers that innervate an inner hair cell (or a group of inner hair cells with similar CF), *N* such processes were run in parallel and their spike times merged. This assumes that subsequent neural stages have access to such a compound spike train (ignoring for simplicity the distinction between high and low spontaneous rate fibers). Optionally, a Gaussian-distributed jitter with standard deviation *σ* can be added to each spike time to approximate synchrony roll-off or other sources of temporal imprecision.

Spike times (produced with arbitrary temporal precision) were converted to a pulse train (100 kHz sampling rate) that was split into a “direct pathwhay” convolved with a decaying exponential kernel modeling an EPSP, and a “delayed pathway” (delay *T*) convolved by a second kernel modeling an IPSP, with time constants *τ_e_* and *τ_i_* respectively. The delay *T* was set to the period of the lower tone. Direct and delayed signals were added (with negative sign for the IPSP-convolved signal), and the sum was half-wave rectified and averaged over the duration of the stimulus, resulting in a single number representing the “activation” of the gating neuron. To simulate the decision made by a subject on every trial (160 trials per condition), the activation for the mistuned pair was compared with that for the in-tune pair, and the decision deemed correct if the former was greater than the latter. This simulation was repeated 1000 times for each condition.

The results presented here are for one channel (haircell), *R*=134 spikes/s, *N* =10 fibers per hair cell, *σ*=0 (no added jitter), *τ_e_*=100 *μs*, *τ_i_*=150 *μs*. Figure 6 compares percentage correct for the model (left column) and the three subjects of Demany et al. (1991) (rightmost columns). Open symbols are for upward mistunings, closed symbols for downward mistunings, and each row is for a different lower frequency. Error bars are ± one standard deviation over repetions divided by the square root of 160 (number of trials in the original study, to gauge the portion of behavioral response variability that might be due to stochastic spike generation).

**FIG. 6.**
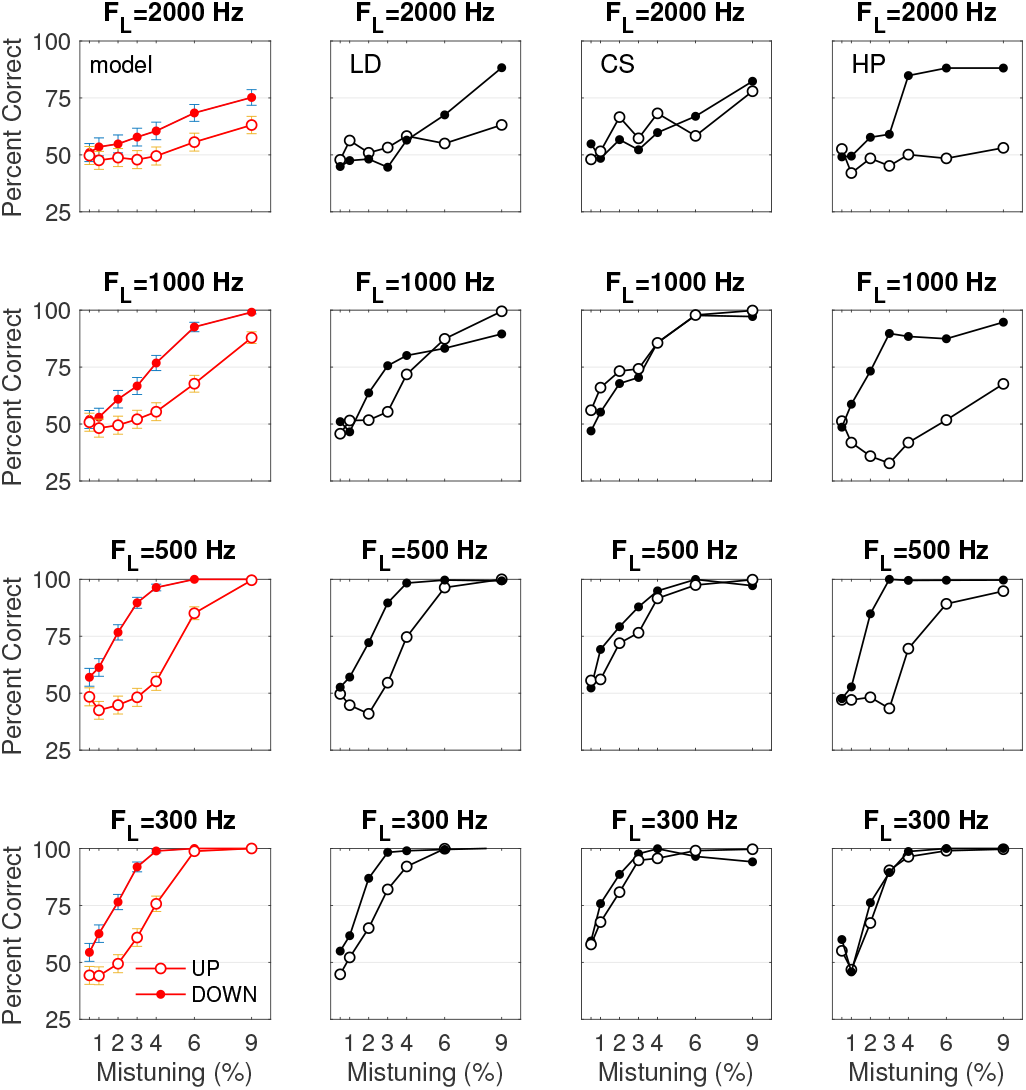
Model simulated data (left column) and behavioral data replotted from Demany et al. (1991) for three subjects (right columns). Filled symbols are for downward mistunings and open symbols upward mistunings. Each row is for a different low frequency, as indicated. Error bars in the left column represent ± the simulated standard deviation divided by the square root of 160 (number of trials in the behavioral study).

Several points are worth noting. First, the model reproduces listeners’ sensitivity to mistuning: detection is near chance for small mistunings and near-perfect for large mistunings, with slopes roughly similar to the behavioral data, shallower at high than low frequencies. Second, sensitivity is greater for negative than positive mistunings, with a hint of bias at small positive mistunings (scores reliably below 50 %). Third, asymmetry is seen at all frequencies, as observed for some (albeit not all) subjects.

Between-subject differences, seen also in other studies (see Discussion), cannot be predicted with a single set of parameters (error bars are too small), but can be reproduced if parameters are allowed to vary between subjects. The asymmetry can be abolished or reversed (as observed for some subjects/conditions) by making the delayed pathway time constant equal to, or shorter than, that of the direct pathway (not shown). Increasing both time constants reduces asymmetry at high frequencies (as observed for subjects LD and CS), decreasing them reduces asymmetry at low frequencies (not shown). A non-zero value of *σ* also abolishes mistuning sensitivity at higher frequencies (not shown). Increasing *N* (e.g. fibers from several inner hair cells pooled) increases mistuning sensitivity, reducing *N* (as might occur due to deafferentation) reduces mistuning sensitivity (not shown). The simulation is based on a single peripheral channel, drawing from a wider range (c.f. Fig. 1) would further increase mistuning sensitivity (not shown), and so-on.

This flexibility is fortunate in that it allows a wide range of observed patterns to be accounted for. It is unfortunate in that the model is difficultly falsifiable, which is objectionable from a Popperian perspective. Better knowledge about the physiology (see Discussion) might allow tighter predictions, for now the values plotted in Fig. 6 carry limited predictive weight.

## III. DISCUSSION

Two things need explaining. The first is *octave affinity*, for both harmonic and melodic octaves. The second is *asymmetric sensitivity to mistuning*. This paper attempts to explain both, but the explanation raises some issues that are discussed here. Anticipating, evidence from physiology leads us to expect an asymmetry, but only weakly predicts its sign. The explanation is directly applicable to the harmonic octave, but accounting for the melodic octave requires additional assumptions.

### a. Affinity or Equivalence?

The octave has been recognized as remarkable since antiquity, and it plays an important role in the music of most cultures (Burns and Ward, 1999; Shepard, 1982). Tones separated by an octave are treated as equivalent for the definition of a chroma class (Bachem, 1950; Burns and Ward, 1999; Deutsch, 1969; Harrison and Pearce, 2020). For two complex tones separated by an octave, all the harmonics of the higher tone (including its fundamental) are also harmonics of the lower tone. Likewise, subharmonics of the lower tone are subharmonics of the higher tone. The lower tone thus “contains” the octave tone, but the converse is not true: octave similarity thus lacks the symmetry property that would be required for “equivalence” in a mathematical sense.

### b. Harmonic vs Melodic Octave

Affinity has been observed for both concurrent sounds (harmonic octave) and sequential sounds (melodic octave). For pure tones, the melodic octave is somewhat elusive as discussed by Demany et al. (2021). While some studies suggest perceptual similarity or even equivalence (Demany and Armand, 1984), others found little evidence that melodic octave intervals are special (Kallman, 1982). An interpretation is that two perceptual qualities are involved, one salient for all listeners (difference in tone height) and the other more subtle and requiring musical literacy (chroma) (Bonnard et al., 2013). Tasks that probe the latter require ignoring the former.

Parsimony lobbies for a common basis for the harmonic and melodic octave, but experimental support has been ambiguous. Of three careful studies by the same group, two concluded that there are distinct mechanisms (Bonnard et al., 2016, 2013), and the third that there is a common mechanism (Demany et al., 2021), see also Demany and Semal (1988, 1990); Demany et al. (1991). For a harmonic octave, mistuning is harder to detect if the higher tone exceeds ~2 kHz (Demany et al., 2021), which is possibly related to the finding of Hartmann et al. (1990) that subjects cannot detect mistuning of a partial of a harmonic complex beyond ~2 kHz. For a melodic octave, mistuning is detectable up to ~4 kHz or higher (Ward, 1954), however, mistuning is easier to detect for harmonic than melodic octaves in the low frequency region (Demany et al., 2021).

### c. Detecting mistuning

For a *harmonic* octave, detection is straightforward. Any mechanism that can detect inharmonicity (in the frequency domain) or aperiodicity (in the time domain) can detect octave mistuning. The model presented in this paper uses a cancellation filter for that purpose. For the *melodic* octave, the situation is more subtle because memory is involved: a trace of the first tone must be retained for comparison with the second tone. The trace of the first tone (which might be the lower or the higher) can be *compared* to a similar trace derived from the second tone, or instead used as a *parameter of a mistuning detector* applied to the second tone (as in this paper).

Terhardt (1974) assumed that every harmonic of a complex tone evokes a “spectral pitch” which is transformed into a series of “virtual pitch” cues via a harmonic template. Cues from all harmonics are then tallied in a weighted histogram (see also Schroeder, 1968) that shows a series of peaks, the largest of which signals the pitch. However, there are other peaks, and Terhardt noted that the largest among them tend to be separated by an octave interval. The trace, in this account, is the subharmonic histogram of the first tone, compared to the subharmonic histogram of the second tone to detect an octave relation (or mistuning thereof).

This explanation relies on the regular pattern of *harmonics* (overtones) of most musical sounds. Their perceptual reality was noted by Oresme in the XIVth century (Bod, 2014), and their physical reality worked out by a line of thinkers from Mersenne in the XVIIth up to Fourier in the XIXth century (de Cheveigné, 2005). Interestingly, the mathematician Fermat, writing to Mersenne in 1636, raised the possibility that the “true” octave might differ from a ratio 2:1 (https://www.musicologie.org/Biographies/m/mersenne.html). The implication of harmonics in *octave affinity* is attributed to Sauveur and Rameau in the 18th century (Christensen, 1987), and *learning* of a template from lifelong exposure to harmonic sounds, notably speech (Terhardt, 1974), to Jean Jacques Dortous de Mairan, contemporary of Rameau (Christensen, 1987).

Instead of harmonics, one can invoke *subharmonics* (super-periods). Ohgushi (1983) invoked for this purpose the first-order interspike interval (ISI) histogram of auditory nerve fiber discharge patterns, that typically shows a series of modes near period multiples. The ISI histogram for the first tone is compared to a similar histogram from the second tone. At the octave, some peaks of the histogram for the higher tone coincide with peaks for the lower tone, hence affinity. A similar idea has been proposed for other auditory nerve fiber interval statistics such as autocorrelation (Cariani and Delgutte, 1996; Tramo, 2005). The principle has much earlier roots in the “coincidence theory of consonance” elaborated in the 16th and 17th centuries (Christensen, 1993; Green, 1969), and indeed in the association of consonant intervals with small integer ratios usually attributed to Pythagoras.

The model proposed in this paper is of the “subharmonic” flavor. The trace, in this case, is the delay parameter *T* of the first filter, conserved and used to tune a new filter applied to the second tone (or the filter itself, “frozen” until the second tone). This sequential operation is straightforward in the case of an *ascending* octave, because the filter fit to the lower tone also cancels the higher, but for a descending octave the filter must be fit to *twice* the period of the first tone. This is less straightforward, but perhaps a skill that a musically-inclined subject might have learned to master. Incidentally, data specific to the descending melodic octave are somewhat sparse. Bonnard et al. (2013); Cuddy and Dobbins (1988) presented notes only in ascending order, while Demany et al. (2021); Demany and Semal (1990); Ward (1954) presented notes in alternation, so ascending and descending intervals contributed to each judgement. However, Bonnard et al. (2016) presented half their subjects with ascending intervals and the other half with descending intervals, and found no reliable difference, consistent with reports of Hartmann (1993) and Rosner (1999). Luo et al. (2014) observed that musical intervals are generally less accurately ranked when descending than ascending (but they did not test octaves), and Vos and Troost (1989) analyzed two music databases and found that descending octaves were half as common as ascending.

### d. Octave stretch and mistuning asymmetry

Octave stretch (preference for a frequency ratio larger than 2) and mistuning asymmetry (greater tolerance for positive than negative mistuning) are distinct phenomena, but that could conceivably be related: in the presence of jitter or noise, stretching the octave might be a “safe” strategy to reduce the likelihood of a perceptually noticeable mistuning. If such is the case, a model of mistuning asymmetry could account for both phenomena, and data related to both would be relevant to constrain that model.

Mistunings are easier to detect for positive than negative deviations for both the harmonic octave (Bonnard et al., 2013; Demany et al., 2021; Demany and Semal, 1988; Demany et al., 1991) and the melodic octave (Demany et al., 2021; Demany and Semal, 1990; Dobbins and Cuddy, 1982). It is however unclear that this is specific to the octave, as a similar stretch has been reported for intervals of minor sixth or greater, whereas smaller intervals are compressed (Burns and Ward, 1999; Rakowski, 1990). The melodic octave is stretched for pure tones (Hartmann, 1993; Ohgushi, 1983) as well as complex tones and sounds of musical instruments (Jaatinen et al., 2019; Sundberg and Lindqvist, 1973).

The melodic stretch is typically a few percent, so a subject must be sensitive to pitch differences of that order, as is the case for most musicians but not all non-musicians (Micheyl et al., 2006). For at least some subjects, the standard deviation of octave matches was indeed smaller than the magnitude of the stretch (Demany and Semal, 1990). The magnitude of the stretch varies markedly across subjects (Demany et al., 2021; Demany and Semal, 1990; Hartmann, 1993; Ward, 1954), and for some subjects, in some conditions, it is close to zero or even *reversed* (Bonnard et al., 2013; Hartmann, 1993; Rosner, 1999). This is an extra burden for a model as it must explain why the phenomenon is observed in some subjects and not others. Large inter-subject differences were also observed for pitch discrimination (Micheyl et al., 2006), and sensitivity to consonance (McDermott et al., 2010).

The stretch is typically larger at higher frequencies than low, at least for the melodic octave (Demany and Semal, 1990; Jaatinen et al., 2019; Ohgushi, 1983). For a harmonic octave, as mentioned earlier, the ability to notice mistuning itself is reduced in the upper range. Bonnard et al. (2013) found that sensitivity to mistuning of either direction improved with level, which they attributed to the increased salience of beats, however, the asymmetry vanished at higher levels. For the melodic octave, Sundberg and Lindqvist (1973) found a complex dependency of asymmetry on subject, frequency, tone nature (pure or complex), and intensity of both tones.

Bonnard et al. (2013) measured discrimination between various frequency ratios in the vicinity of 2:1 for concurrent pure tone pairs. Discrimination from a reference ratio of 1.96:1 was unexpectedly better for a test ratio of 2:1 than for a larger test ratio. From this, they concluded that the true octave ratio (2:1) is “special”. Apparently, the subjective versus objective mistuning function is “V”-shaped with a minimum at 2:1, but with a steeper slope on the lower (<1 octave) than higher (>1 octave) side, consistent with Fig. 5 (bottom right). Bonnard et al. (2016) also found that perceptual fusion was maximal for a ratio close to 2:1, decreasing more rapidly for downward than upward mistunings, which again suggests an asymmetric tolerance for mistuning from the optimal ratio 2:1, rather an optimal ratio larger than 2:1. The asymmetry was not observed for mildly hearing-impaired subjects (Bonnard et al., 2017).

It has also been suggested that the asymmetry might reflect an *aesthetic* bias (e.g. Demany et al., 2021), which begs the question of what drives the aesthetics. A possibility is that mistuning is unpleasant, and the bias results from upwards mistunings being less noticeable, but the question then is why mistuning is unpleasant. A speculative explanation is that harmonicity-based segregation mechanisms help maintain the clarity of a complex musical scene, pleasurable according to the concept of *perceptual fluency* (Reber et al., 1998). If so,the aesthetics themselves would be an emergent property.

### e. Physiological Basis

Excitatory-inhibitory (EI) interaction is found throughout the nervous system. High-bandwidth auditory representations are supported by temporally-specialized neural circuits, some excitatory and others inhibitory (Joris and van der Heijden, 2019; Stange-Marten et al., 2017). Fast EI interaction has been identified as early as the cochlear nucleus (Campagnola and Manis, 2014; Joris and Smith, 1998; Kuenzel, 2019; Ngodup et al., 2020; Paolini et al., 2004; Xie and Manis, 2013) and as late as the the dendritic fields of the inferior colliculus (Caspari et al., 2015; Chen et al., 2019), in addition to intermediate stages within the superior olivary complex (Franken et al., 2018; Grothe, 2003; Myoga et al., 2014; Roberts et al., 2013) and nuclei of lateral lemniscus (Caspari et al., 2015).

In some instances, EI interaction has been attributed to binaural processing, for example estimation of interaural level differences (ILD) (Tollin and Yin, 2005), detection of onset timing differences (Franken et al., 2018), or sharpening of sensitivity to interaural time difference (ITD) (Myoga et al., 2014). In others, the processing seems to be monaural. For example, neurons in the ventral nucleus of the lateral lemniscus (VNLL) receive, via calyx-like synapses, excitatory input from contralateral cochlear nucleus (CN) and ipsilateral medial nucleus of the trapezoid body (MNTB) (Caspari et al., 2015), neurons in superior periolivary nucleus (SPON) and VNLL receive input from contralateral octopus cells and ipsilateral MNTB (Felix II et al., 2017). Cells in MSO also receive EI input and a certain number of them are monaural (Grothe, 2000).

Inhibition requires an extra synapse and thus might be expected to lag excitation. This has indeed been observed (Xie and Manis, 2013), but the opposite has also been found (Nayagam et al., 2005; Paolini et al., 2004; Roberts et al., 2013), presumably reflecting properties of the inhibitory pathways that favor transmission speed, such as thick myelinated axons or secure synapses (Grothe, 2003; Stange-Marten et al., 2017).

The dynamics of interaction are usually thought to be governed by the time constants of excitatory and inhibitory post synaptic events, for example via a subtractive mechanism analogous to Fig. 4 (right) (Roberts et al., 2013; van der Heijden et al., 2013). Inhibitory events (IPSPs or IPSCs, inhibitory post-synaptic currents) are often reported to be more sluggish than excitatory (EPSPs or EPSCs, excitatory post-synaptic currents) (Campagnola and Manis, 2014; Caspari et al., 2015). Examples of decay time constants reported for inhibition and excitation, respectively, are ~10 ms vs ~ 0.5 ms in mouse AVCN bushy cells (Campagnola and Manis, 2014), ~2.5 ms vs ~0.3 ms in mouse ventrolateral VNLL (Caspari et al., 2015), or ~1.5 ms vs 0.3 ms in mouse MSO (Myoga et al., 2014), etc. In contrast, Pilati et al. (2016) reported closely matched time constants of EPSCs and IPSCs in mouse LSO.

However, the dynamics of EI interaction might not be directly governed by the duration of post-synaptic current or potential events: Franken et al. (2021) observed in LSO a window of binaural interaction much shorter (~0.5 ms or less) than the IPSP decay time constant (~5 ms), and with remarkably steep slopes (transition from maximum to suppressed firing on the order of 0.1 ms). Thus, the relatively short time constants (100 and 150 μs) chosen for the simulation (section Methods and Results) are not entirely implausible.

In summary, there is ample evidence that dynamics of excitation and inhibition might differ between neural pathways, some delayed relative to others, as required by the account proposed in this paper. However, the evidence is sufficiently complex and diverse to support either direction of octave shift. Slower dynamics for the delayed pathway should lead to a positive shift (consistent with behaviour), whereas faster dynamics would lead to the opposite shift. Better knowledge about the locus of interaction and its properties is required for tighter predictions.

### f. Other Explanations

A simple explanation is a response or classification bias due to the structure of the response or category set. If responses are limited to chroma within the octave, the listener might maintain a sharp boundary between the major seventh and the octave, but not necessarily between the octave and the minor ninth. A similar hypothesis could explain why the perceptual boundary between minor and major second is low (Rakowski, 1990; Vurma and Ross, 2006), and more generally the tendency for large intervals (not just the octave) to stretch and small intervals to shrink. This explanation seems less likely in a non-musical psychoacoustic setting, such as discriminating mistuned from in-tune intervals (Bonnard et al., 2013).

Another proposal, also rooted in music, relates octave enlargement to the “Pythagorean comma”, which is the frequency ratio between twelve fifths (1.5^12^) and seven octaves, (2^7^), which amounts to roughly 3.4 cents per octave (Hubbard, 2021). This is in the ballpark of some perceptual shifts, but it is not clear why they would not then be equally distributed along the seven octaves, rather than the stretch being small (even negative) at low frequencies and large (up to 50 cents) at high frequencies. Nor is it clear how shifts could be subject-dependent, or level-dependent. A different, physics-based explanation is that the scale of notes on the piano is stretched to reduce the perceptual effects of beats that result from the inharmonic spectrum of strings (Giordano, 2015). The piano is often used as a reference to tune other instruments, and it is conceivable that this led musicians to expect octaves to be mistuned. The so-called “Railback” tuning curve is steep at the low- and high-frequency ends, consistent with a larger octave stretch in the high register but not with the small or negative octave stretch observed in the low register.

As mentioned earlier, Terhardt (1974) proposed that mutual masking between partials shifts their “spectral pitch”, resulting in a shift of the “virtual pitch” cues that they evoke via a harmonic template, and a mismatch when cues for the lower and higher tone are lined up. The same effects could also affect learning of that template. The explanation is weakened by the failure of Peters et al. (1983) to replicate the shifts upon which this model is predicated.

Also mentioned earlier, Ohgushi (1983) explained octave shifts on the basis of the first-order interspike interval (ISI) histogram of auditory nerve fiber discharges (e.g. Rose et al., 1967). His explanation exploits two quirks of first-order ISIs: (a) Spikes cannot occur at intervals shorter than the *absolute refractory period* of the auditory nerve (~1 ms), and their probability remains depressed over the next ~5 ms (relative refractory period) so the histogram is truncated near its origin and its first mode or modes may be missing or shifted rightward. (b) At low frequency and/or high discharge rate, multiple spikes may occur within a period, in which case intervals that span periods are shortened, leading to a leftward shift of the histogram modes. Leftward shifts are more common for low frequencies, rightward shifts more common for higher frequencies, hence a mismatch when histograms for octave-spaced tones are lined up, reduced by increasing the frequency of the higher tone.

This explanation is not entirely satisfactory. The first-order ISI histogram was used in early studies for technical reasons, but it does not well represent the information carried by the spike train. More recent studies prefer the all-order ISI histogram (autocorrelation) (e.g. Cariani and Delgutte, 1996) and shuffled versions thereof (Joris et al., 2006, 2003). A first issue is that the first-order ISI histogram is relatively noisy because a *N*-spike train offers only *N* – 1 intervals, in contrast to the ~ *N*^2^/2 all-order intervals exploitable by the all-order ISI histogram, or the ~ (*KN*)^2^/2 all-order intervals within a group of *K* fibers as reflected by a shuffled all-order histogram (Joris et al., 2006, 2003)). A second issue is that the shape of the first-order histogram changes drastically with discharge rate (and thus level): a high rate induces multiple spikes within the same period, leading to an exagerated zero-order mode and a leftward shift of subsequent modes (this is the quirk (b) mentioned above). These reasons make the first-order ISI histogram an implausible basis for a model.

Hartmann (1993) and McKinney and Delgutte (1999). McKinney and Delgutte (1999) adapted Ohgushi’s model to the more plausible all-order ISI histogram. McKinney and Delgutte (1999) recorded from a large number of auditory nerve fiber recordings in the cat and developed a quantitative model of melodic octave shifts. The model is highly constrained by the physiological data, with only one free parameter, an advantage in terms of falsifiability. However, this feature also makes it harder to account for inter-individual differences. Furthermore, the model predicts only small asymmetries at low frequencies (unless the implausible first-order ISI histogram is used), and requires relatively long delays (on the order of 5 periods). Perhaps the most serious weakness is that the refractoriness-based property that it relies on is lost when spikes from multiple fibers (e.g. from the same haircell) are pooled. Forbidding such pooling means assuming that each neuron contributing to calculate the histogram receives input from at most one fiber, wasting much of the information. This is a strong assumption.

To summarize, multiple explanations of octave stretching can be envisaged, in addition to that proposed in this paper. Each is deficient in certain aspects, so we cannot say, at this point, that the problem is solved. Hopefully, the present model is a step in the right direction.

### g. Limitations

A first weakness of the model is that, while it accounts for the possibility of a systematic mistuning of the octave, it only weakly predicts its magnitude or sign. To do so would require identifying the locus of its implementation within the auditory system, and verifying that the time constants agree with the assumptions of the model.

A second weakness is that the model is not immediately applicable to the melodic octave. To account for mistuning asymmetry of both ascending and descending octaves, effects must apply equally to the first tone (when the trace is established) and at the second tone (where it is applied or compared with a trace derived from the second tone), but with a greater impact on high than low frequency tones. The details of how this would happen have not been worked out.

### h. Significance

Keeping limitations of the model in mind, we can revisit the claim, made earlier, that the careful observation of a relatively subtle effect may yield wider insights into hearing. The logic is that, since the effect is reliably established, and cannot easily be explained otherwise, each of the model’s assumptions receives some degree of support. Reviewing:

- Pitch perception and and harmonic fusion involve processing of time domain patterns in the auditory brain, in contrast to a purely spectral account (e.g. Terhardt, 1974)
- Processing involves a spike coincidence or anticoincidence counting process akin to autocorrelation or cancellation (e.g Cariani and Delgutte, 1996; de Cheveigné, 1998; Meddis and Hewitt, 1991) in contrast, for example, to first-order interspike interval statistics (Ohgushi, 1983).
- Cancellation is favored over correlation as it can help ensure *invariance* to competing periodic sounds (de Cheveigné, 2021), suggesting that asymmetry is an emergent property of a mechanism that addresses wider needs. Cancellation can be implemented on the basis of correlation (in this sense the two are equivalent) but this entails an additional implementation complexity (de Cheveigné, 2021).
- Cancellation (and correlation) may be affected by details of their neural implementation, such as the shape of PSPs involved in coincidence detection.

Neural cancellation was previously effective in a model of pitch shifts of mistuned partials (de Cheveigné, 1999; Hartmann and Doty, 1996; Hartmann et al., 1990). The asymmetric shape of PSPs was also invoked in a model to explain why the pitch of a peak in frequency modulation is more salient than that of a trough (de Cheveigné, 2000; Demany and McAnally, 1994), although that model involved autocorrelation rather than cancellation, and the asymmetry resulted from the asymmetric shape of PSPs rather than different time constants along different pathways.

To the extent that octave mistuning asymmetry is indeed explanable with these assumptions, it may offer wider insights as to how hearing works.

## CONCLUSION

The asymmetry of mistuning of the harmonic octave can be explained by a model of excitatory-inhibitory interaction within the auditory brain. A difference in time constants of excitation and inhibition causes the neural circuit to be more sensitive to a negative than a positive mistuning of the higher tone. This paper reviews the phenomenon, describes the model, reviews relevant data from auditory physiology, and discusses evidence for, and against, the model.

## ACKNOWLEDGMENTS

This work was supported by grants ANR-10-LABX-0087 IEC, ANR-10-IDEX-0001-02 PSL, and ANR-17-EURE-0017. Laurent Demany contributed many ideas and useful criticism to earlier versions of this paper, which also benefitted from useful comments from the Editor (Philip Joris) and two anonymous reviewers.

## References

Bachem A (1950) Tone height and tone chroma as two different pitch qualities. Acta Psychologica 7:80–8.

Bod R (2014) A new history of the humanities: the search for principles and patterns from antiquity to the present. Oxford Scholarship Online 52:52–0622–52–0622.

Bonnard D, Dauman R, Semal C, Demany L (2016) Harmonic fusion and pitch affinity: Is there a direct link? Hearing Research 333:247–254.

Bonnard D, Dauman R, Semal C, Demany L (2017) The Effect of Cochlear Damage on the Sensitivity to Harmonicity Ear & Hearing 28 247–254.

Bonnard D, Micheyl C, Semal C, Dauman R, Demany L (2013) Auditory discrimination of frequency ratios: The octave singularity. Journal of Experimental Psychology: Human Perception and Performance 39:788–801.

Burns E, Ward W (1999) Intervals, scales, and tuning In Deutsch D, editor, The Psychology of Music, pp. 215–264. Academic Press.

Campagnola L, Manis PB (2014) A Map of Functional Synaptic Connectivity in the Mouse Anteroventral Cochlear Nucleus. Journal of Neuroscience 34:2214–2230.

Cariani P, Delgutte B (1996) Neural correlates of the pitch of complex tones. i. pitch and pitch salience. J. Neurophysiol. 76:1698–1716.

Caspari F, Baumann VJ, Garcia-Pino E, Koch U (2015) Heterogeneity of Intrinsic and Synaptic Properties of Neurons in the Ventral and Dorsal Parts of the Ventral Nucleus of the Lateral Lemniscus. Frontiers in Neural Circuits 9.

Chen C, Read HL, Escabí MA (2019) A temporal integration mechanism enhances frequency selectivity of broadband inputs to inferior colliculus. PLOS Biology 17:e2005861.

Christensen T (1987) Eighteenth-Century Science and the “Corps Sonore:” The Scientific Background to Rameau’s ‘‘Principle of Harmony”. Journal of Music Theory 31:23.

Christensen T (1993) Rameau and musical thought in the enlightenment Cambridge University Press.

Cuddy LL, Dobbins PA (1988) Octave Discrimination: Temporal and Contextual Effects. Canadian Acoustics 16:3–13.

de Cheveigné A (1985) A nerve fiber discharge model for the study of pitch. Transactions of the Committee on Speech Research/Hearing Research !The Acoustical Society of Japan, Tokyo S85-37:279–286.

de Cheveigné A (1998) Cancellation model of pitch perception. J Acoust Soc Am 103:1261–1271.

de Cheveigné A (2000) A model of the perceptual asymmetry between peaks and troughs of frequency modulation. J. Acoust. Soc. Am 107:2645–2656.

de Cheveigné A (2005) Pitch perception models In C P, A O, editors, Pitch - Neural coding and perception, pp. 169–233. Springer.

de Cheveigné A (2021) Harmonic cancellation - a fundamental of auditory scene analysis. Trends in Hearing in press.

de Cheveigné A (1999) Pitch shifts of mistuned partials: A time-domain model. The Journal of the Acoustical Society of America 106:887–897.

Demany L, McAnally KI (1994) The perception of frequency peaks and troughs in wide frequency modulations. J. Acoust. Soc. Am. 96:706–715.

Demany L, Armand F (1984) The perceptual reality of tone chroma in early infancy. J. Acoust. Soc. Am. 76:57–66.

Demany L, Monteiro G, Semal C, Shamma S, Carlyon RP (2021) The perception of octave pitch affinity and harmonic fusion have a common origin. Hearing Research 404:108213.

Demany L, Semal C (1988) Dichotic fusion of two tones one octave apart: Evidence for internal octave templates. The Journal of the Acoustical Society of America 83:687–695.

Demany L, Semal C (1990) Harmonic and melodic octave templates. The Journal of the Acoustical Society of America 88:2126–2135.

Demany L, Semal C, Carlyon RP (1991) On the perceptual limits of octave harmony and their origin. The Journal of the Acoustical Society of America 90:3019–3027.

Deutsch D (1969) Music recognition. Psychological Review 76:300–307.

Dobbins PA, Cuddy LL (1982) Octave discrimination: An experimental confirmation of the “stretched” subjective octave. The Journal of the Acoustical Society of America 72:411–415.

Felix II RA, Gourévitch B, Gómez-Álvarez M, Leijon SCM, Saldaña E, Magnusson AK (2017) Octopus Cells in the Posteroventral Cochlear Nucleus Provide the Main Excitatory Input to the Superior Paraolivary Nucleus. Frontiers in Neural Circuits 11:37.

Franken TP, Bondy BJ, Haimes DB, Goldwyn JH, Golding NL, Smith PH, Joris PX (2021) Glycinergic axonal inhibition subserves acute spatial sensitivity to sudden increases in sound intensity. eLife 10:e62183.

Franken TP, Joris PX, Smith PH (2018) Principal cells of the brainstem’s interaural sound level detector are temporal differentiators rather than integrators. eLife 7:e33854.

Giordano N (2015) Explaining the Railsback stretch in terms of the inharmonicity of piano tones and sensory dissonance. The Journal of the Acoustical Society of America 138:2359–2366.

Green BL (1969) The harmonic series from Mersenne to Rameau: An historical study of circumstances leading to its recognition and application to music Ph.D. diss., Ohio State University.

Grothe B (2000) The function of the medial superior olive in small mammals: temporal receptive fields in auditory analysis. Journal of Comparative Physiology A 186:413–423.

Grothe B (2003) New roles for synaptic inhibition in sound localization. Nat Rev Neu-rosci 4:413–423.

Harrison PMC, Pearce MT (2020) Simultaneous consonance in music perception and composition. Psychological Review 127:540–550,). https://doi.org/10.1038/nrn1136.

Hartmann WM (1993) On the origin of the enlarged melodic octave. The Journal of the Acoustical Society of America 93:3400–3409.

Hartmann WM, Doty SL (1996) On the pitches of the components of a complex tone. The Journal of the Acoustical Society of America 99:567–578.

Hartmann W, McAdams S, Smith BK (1990) Hearing a mistuned harmonic in an otherwise periodic complex tone. J. Acoust. Soc. Am. 88:1712–1724.

Hubbard TL (2021) The Pythagorean comma and preference for a stretched octave. Psychology of Music p. 030573562110089.

Jaatinen J, P’átynen J, Alho K (2019) Octave stretching phenomenon with complex tones of orchestral instruments. The Journal of the Acoustical Society of America 146:3203–3214.

Joris PX, Louage DH, Cardoen L, van der Heijden M (2006) Correlation Index: A new metric to quantify temporal coding. Hearing Research 216-217:19–30.

Joris PX, (2003) Interaural time sensitivity dominated by cochlea-induced envelope patterns. J Neurosc 23:6345–6350.

Joris PX, Smith PH (1998) Temporal and Binaural Properties in Dorsal Cochlear Nucleus and Its Output Tract. The Journal of Neuroscience 18:10157–10170.

Joris PX, van der Heijden M (2019) Early Binaural Hearing: The Comparison of Temporal Differences at the Two Ears. Annual Review of Neuroscience 42:433–457.

Kallman HJ (1982) Octave equivalence as measured by similarity ratings. Perception & Psychophysics 32:37–49.

Kuenzel T (2019) Modulatory influences on time-coding neurons in the ventral cochlear nucleus. Hearing Research 384:107824.

Lewis PAW, Shedler GS (1979) Simulation of nonhomogeneous Poisson processes by thinning. Naval Res. Logistics Quart. 26:403–413.

Licklider J (1951) A duplex theory of pitch perception. Experientia 7:128–134.

Lu Y, Liu Y, Curry RJ (2018) Activity-dependent synaptic integration and modulation of bilateral excitatory inputs in an auditory coincidence detection circuit: Synaptic integration and modulation in nucleus laminaris. The Journal of Physiology 596:1981–1997.

Luo X, Masterson ME, Wu CC (2014) Melodic interval perception by normal-hearing listeners and cochlear implant users. The Journal of the Acoustical Society of America 136:1831–1844.

McDermott JH, Lehr AJ, Oxenham AJ (2010) Individual Differences Reveal the Basis of Consonance. Current Biology 20:1035–1041.

McKinney MF, Delgutte B (1999) A possible neurophysiological basis of the octave enlargement effect. The Journal of the Acoustical Society of America 106:2679–2692.

Meddis R, Hewitt MJ (1991) Virtual pitch and phase sensitivity of a computer model of the auditory periphery. I: Pitch identification. The Journal of the Acoustical Society of America 89:2866–2882.

Micheyl C, Delhommeau K, Perrot X, Oxenham AJ (2006) Influence of musical and psychoacoustical training on pitch discrimination. Hearing Research 219:36–47.

Moore B, Glasberg B (1983) Suggested formulae for calculating auditory-filter bandwidths and excitation patterns. J. Acoust. Soc. Am. 74:750–753.

Moore BCJ, Peters RW, Glasberg BR (1985) Thresholds for the detection of inharmonicity in complex tones. The Journal of the Acoustical Society of America 77:1861–1867.

Myoga MH, Lehnert S, Leibold C, Felmy F, Grothe B (2014) Glycinergic inhibition tunes coincidence detection in the auditory brainstem. Nature Communications 5:3790.

Nayagam DAX, Clarey JC, Paolini AG (2005) Powerful, Onset Inhibition in the Ventral Nucleus of the Lateral Lemniscus. Journal of Neurophysiology 94:1651–1654.

Ngodup T, Romero GE, Trussell LO (2020) Identification of an inhibitory neuron subtype, the l-stellate cell of the cochlear nucleus. eLife 9:e54350.

Ohgushi K (1983) The origin of tonality and a possible explanation of the octave enlargement phenomenon. The Journal of the Acoustical Society of America 73:1694–1700.

Paolini AG, Clarey JC, Needham K, Clark GM (2004) Fast Inhibition Alters First Spike Timing in Auditory Brainstem Neurons. Journal of Neurophysiology 92:2615–2621.

Patterson RD (1976) Auditory filter shapes derived with noise stimuli. The Journal of the Acoustical Society of America 59:640–654.

Peters RW, Moore BCJ, Glasberg BR (1983) Pitch of components of complex tones. J. Acoust. Soc. Am. 73:924–929.

Pilati N, Linley DM, Selvaskandan H, Uchitel O, Hennig MH, Kopp-Scheinpflug C, Forsythe ID (2016) Acoustic trauma slows AMPA receptor-mediated EPSCs in the auditory brainstem, reducing GluA4 subunit expression as a mechanism to rescue binaural function: Acoustic trauma slows EPSCs in the LSO. The Journal of Physiology 594:3683–3703.

Rakowski A (1990) Intonation Variants of Musical Intervals in Isolation and in Musical Contexts. Psychology of Music 18:60–72.

Reber R, Winkielman P, Schwarz N (1998) Effects of Perceptual Fluency on Affective Judgments. Psychological Science 9:45–48.

Roberts MT, Seeman SC, Golding NL (2013) A Mechanistic Understanding of the Role of Feedforward Inhibition in the Mammalian Sound Localization Circuitry. Neuron 78:923–935.

Rose JE, Brugge JF, Anderson DJ, Hind JE (1967) Phase-locked response to low-frequency tones in single auditory nerve fibers of the squirrel monkey. Journal of Neurophysiology 30:769–793.

Rosner BS (1999) Stretching and Compression in the Perception of Musical Intervals. Music Perception 17:101–113.

Schroeder M (1968) Period histogram and product spectrum: new methods for fundamentalfrequency measurement. J. Acoust. Soc. Am. 43:829–834.

Shepard R (1982) Geometrical approximations to the structure of musical pitch. Psychological Review 89:305–333.

Slaney M (1993) An efficient implementation of the patterson-holdsworth auditory filter bank technical report 35, Apple Computer.

Stange-Marten A, Nabel AL, Sinclair JL, Fischl M, Alexandrova O, Wohlfrom H, Kopp-Scheinpflug C, Pecka M, Grothe B (2017) Input timing for spatial processing is precisely tuned via constant synaptic delays and myelination patterns in the auditory brainstem. Proceedings of the National Academy of Sciences 114:e4851–E4858.

Sundberg JEF, Lindqvist J (1973) Musical octaves and pitch. The Journal of the Acoustical Society of America 54:922–929.

Terhardt E (1974) Pitch, consonance, and harmony. The Journal of the Acoustical Society of America 55:1061–1069.

Tollin DJ, Yin TCT (2005) Interaural Phase and Level Difference Sensitivity in Low-Frequency Neurons in the Lateral Superior Olive. The Journal of Neuroscience 25:10648–10657.

Tramo MJ (2005) Neurophysiology and neuroanatomy of pitch perception: Auditory cortex. Annals of the New York Academy of Sciences 1060:148–174.

van der Heijden M, Lorteije J, Plau(š)ka A, Roberts M, Golding N, Borst J (2013) Directional Hearing by Linear Summation of Binaural Inputs at the Medial Superior Olive. Neuron 78:936–948.

Vos PG, Troost JM (1989) Ascending and Descending Melodic Intervals: Statistical Findings and Their Perceptual Relevance. Music Perception 6:383–396.

Vurma A, Ross J (2006) Production and Perception of Musical Intervals. Music Perception 23:331–344.

Ward D (1954) Subjective Musical Pitch. J. Acoust. Soc. Am. 26:369–380.

Xie R, Manis PB (2013) Target-Specific IPSC Kinetics Promote Temporal Processing inAuditory Parallel Pathways. Journal of Neuroscience 33:1598–1614.

